# Intraprocedural endothelial cell seeding of arterial stents via biotin/avidin targeting mitigates in-stent restenosis

**DOI:** 10.1101/2022.05.25.493423

**Authors:** Ivan S Alferiev, Bahman Hooshdaran, Benjamin B Pressly, Stanley J Stachelek, Philip W. Zoltick, Michael Chorny, Robert J Levy, Ilia Fishbein

## Abstract

Impaired endothelialization of endovascular stents has been established as a major cause of in-stent restenosis and late stent thrombosis[1]. Attempts to enhance endothelialization of inner stent surfaces by pre-seeding the stents with endothelial cells in vitro prior to implantation are compromised by cell destruction during high-pressure stent deployment. Herein, we report on the novel stent endothelialization strategy of post-deployment seeding of biotin-modified endothelial cells to avidin-functionalized stents. Acquisition of an avidin monolayer on the stent surface was achieved by consecutive treatments of bare metal stents (BMS) with polyallylamine bisphosphonate, an amine-reactive biotinylation reagent and avidin. Biotin-modified endothelial cells retain growth characteristics of normal endothelium and can express reporter transgenes. Under physiological shear conditions, a 50-fold higher number of recirculating biotinylated cells attached to the avidin-modified metal surfaces compared to bare metal counterparts. Delivery of biotinylated endothelial cells to the carotid arterial segment proximal to the implanted avidin-modified stent in rats results in immediate cell binding to the stent struts and is associated with a 30% reduction of in-stent restenosis in comparison with BMS.

## Introduction

Despite remarkable progress in prevention, diagnosis, and treatment, atherosclerosis remains the leading cause of death in industrialized societies [2]. According to a 2019 AHA report, coronary heart disease, peripheral artery disease (PAD) and cerebrovascular disease affect 18.2, 6.8, and 7 million Americans, respectively [2]. Stent angioplasty is the therapy of choice for symptomatic occlusive vascular disease. However, in-stent restenosis (ISR) presents a formidable problem, only partially addressable by drug-eluting stents (DES [3,4]), which perform sub-optimally in patients with renal failure, diabetes, and those with small vessel diameter [5]. Moreover, by inhibiting re-growth of endothelium, DES increase incidence of late stent thrombosis [6]. The economic burden of ISR is at least $2.8 billion a year in the USA alone [7]. Accelerated restoration of a functional endothelial layer in the injured arteries following stent angioplasty has been pursued over the past 25 years as a means of preventing uncontrolled neointimal expansion and subsequent in-stent restenosis (ISR) [8]. To achieve enhanced re-growth of endothelium over the denuded arterial segments, a pre-deployment seeding of stent struts with autologous endothelial cells (EC) [9–16] was previously investigated. These studies established adherence of endothelial cells to the metal substrate and their proliferation under both flow and no flow conditions. Some of these experiments further analyzed the cell coverage of struts before and after *ex vivo* or *in vivo* stent expansion and found that cells growing on the internal (adluminal) stent surface disappear after coming in direct contact with the inflated balloon, while most of the cells grown on the external (abluminal) and the side aspects of the struts endure the impact. The adluminal endothelial cells (EC) demolished during stent deployment represent a natural interface between the implanted stents and circulating blood. In that capacity, adluminal EC play a barrier role protecting the stent surface from the ingress of platelets and inflammatory leukocytes that initiate the chain of events culminating in ISR[1]. Protocols involving stent surface modification to expedite the post-deployment capture and proliferation of circulating endothelial progenitor cells (EPC) have also been explored [17–21]. To date, no major clinical success of either pre-or post-deployment stent endothelialization has been demonstrated [22,23] apparently due to failure to establish integral EC/EPC coverage, which is paramount for aborting platelet attachment to the struts and preventing inflammatory cell recruitment.

This study investigates a novel enhanced endothelialization strategy based on delivering biotin-modified syngeneic EC to luminal aspects of avidin-functionalized stents at the time of stent implantation and their instant tethering to the stent struts via biotin/avidin interactions. This method supports the immediate formation of a functional endothelial layer at the interface between the stent and blood. It provides passivation of the stent-treated vasculature by blocking the ingress of cellular and humoral elements pivotal to neointimal formation and development of ISR.

## Materials and Methods

### Cells

Primary rat aortic endothelial cells (RAEC; Cell Applications, San Diego, CA) were cultured in EGM2 medium (Lonza, Walkersville, MD) and used at passages 4-7. For some experiments the cells were stably transduced with HIV-1-based lentiviral vectors co-encoding eGFP and luciferase reporters (LV_CMV_-eGFP/Luc) at MOI of 10^5^ and further expanded. The lentiviral transfer plasmid, viral packaging, and titering of the viral particles have been previously described[24]. Briefly, using standard molecular biology cloning, eGFP, ribosome skipping Thosia asigna virus 2A, and luc2 (from pGL4.10, Cat. # E6651, Promega, Madison, WI) DNA molecules were inserted and confirmed by Sanger sequencing. The VSVg pseudotyped viral particles were generated in HEK 293T cells by polyethylenimine mediated transient co-transfection. Viral supernatants at 72 hours post-transfection were concentrated by ultracentrifugation.

### Metal surface modification with avidin

316 grade stainless steel stents (Laserage, Waukegan, MI), foil sheet coupons (Goodfellow, Coraopolis, PA) or cylindrical tubing inserts (MicroGroup, Medway, MA) were washed in isopropanol and chloroform at 60°C with shaking for 10 min each, and heated to 200°C for 30 min. The samples were then incubated in 1% polyallylamine bisphosphonate (PAB[25]) at 72°C with shaking for 2 hours, washed in double distilled water (DDW) and reacted with the biotinylation reagent, EZ-Link™ sulfo-NHS-LC-LC-biotin (Thermo Scientific, Waltham, MA) in a carbonate/bicarbonate buffer (pH=9.3) at 4 mM concentration at 37°C with shaking for 1 hour. After thorough DDW and PBS washing, the samples were incubated with 10-50 mg/ml solution of ImmunoPure Avidin (Thermo Scientific, Waltham, MA) in 1% BSA/PBS at 37°C with shaking for 30 min and extensively washed with PBS (Fig. 1, Steps 1-3). In some experiment cell culture plates were coated with type I rat tail collagen (50 µg/ml) overnight. The plates were then treated with EZ-Link™ sulfo-NHS-LC-LC-biotin and avidin as described above for the metal specimens.

**Fig. 1.**
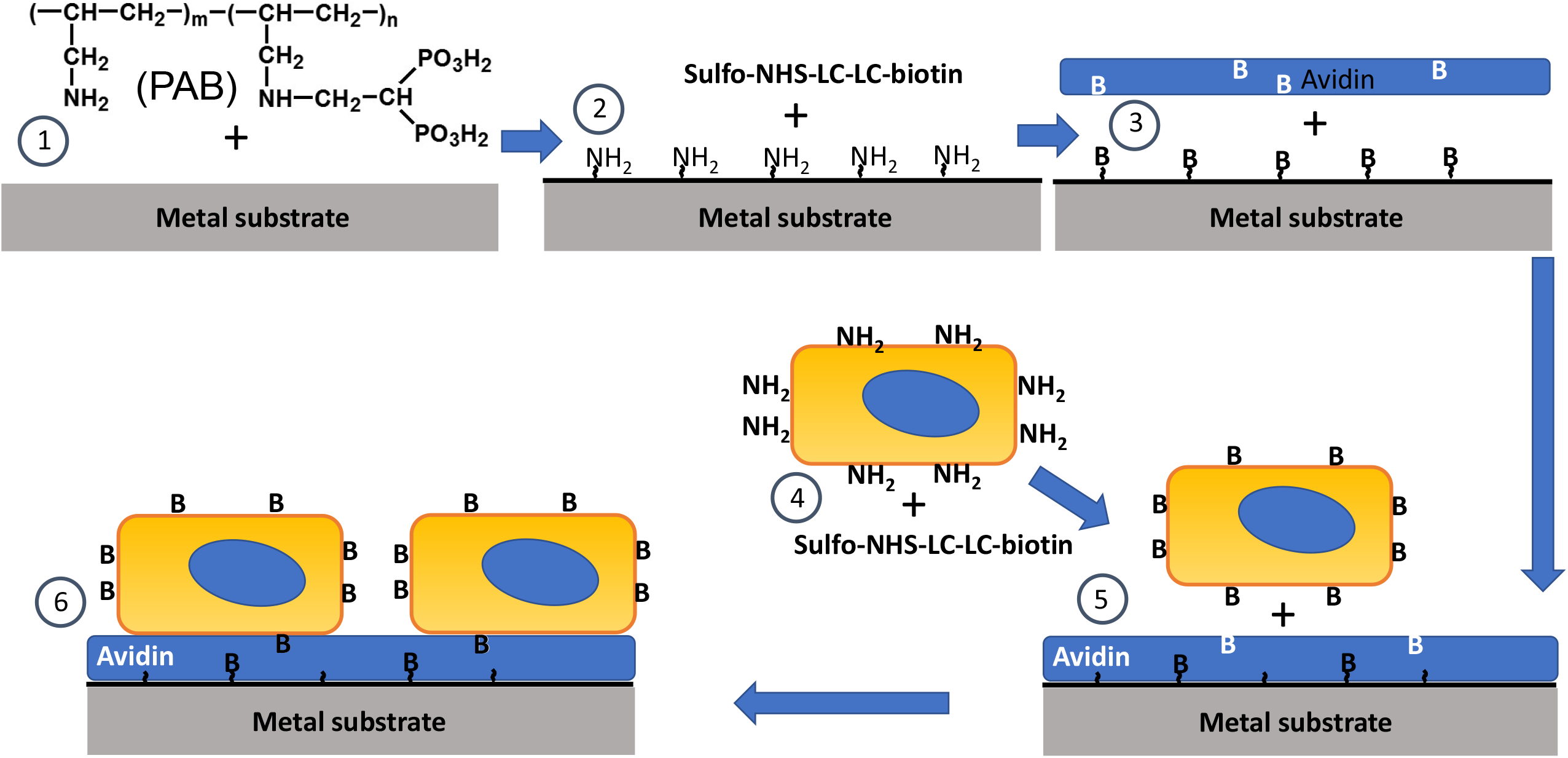
Endothelial cell affinity immobilization on the steel substrate. Step 1: Metal samples are exposed to aqueous polyallylamine bisphosphonate (PAB) solution resulting in permanent surface modification via formation of coordination bonds between the metal atoms and bisphosphonate groups of PAB to exhibit primary amine groups available for chemical conjugation reactions. Step 2: Aminated metal substrate is reacted with an amine-reactive biotinylation agent (sulfo-NHS-LC-LC-biotin). Step 3: Biotinylated metal substrate is exposed to avidin to attain a monolayer of avidin on the metal surface. Step 4: Endothelial cells are surface modified with biotin via the reaction of sulfo-NHS-LC-LC-biotin with amine group of proteins on the cell surface. Step 5: Biotinylated endothelial cells are brought in contact with avidin-modified metal samples resulting in augmented cell binding to the metal substrate (Step 6).

### RAEC surface modification and substrate immobilization

Low passage RAEC were harvested by trypsinization, pelleted by centrifugation, resuspended in 1 ml of 1-4 mM sulfo-NHS-LC-LC-biotin/PBS, placed in a tube mixer and reacted at room temperature for 15 min. Ten ml of complete medium was then added to stop the reaction. The cells were pelleted by centrifugation and washed twice in the medium to remove the excess of sulfo-NHS-LC-LC-biotin. RAEC were then counted using a hemocytometer, and 10_5_-10^6^ cells were placed in individual plastic wells with avidin-modified or control steel samples. Additionally, for control purposes, non-modified RAEC were used when appropriate.

### Avidin depletion assay

Non-modified 1 cm x 1 cm stainless steel foil coupons (n=3) and foils samples modified with PAB and sulfo-NHS-LC-LC-biotin (n=3) were individually incubated with 1 ml of 1 µg/ml solution of Cy3.5-streptavidin (Rockland, Pottstown, PA) in 1% BSA/PBS at 28°C with shaking for 30 min. Additionally 1 ml aliquots (n=3) of the same Cy3.5-streptavidin formulation were processed without metal samples to account for potential adsorption of fluorescently labelled streptavidin on the walls of a glass vial. Residual fluorescence of the solution was then determined using a SpectraMax Gemini EM fluorimeter (Molecular Devices, San Jose, CA) and images of the foil samples were taken. Concentration and immobilization density of streptavidin on the surface of biotin-derivatized steel substrate was calculated using the known initial concentration of Cy3.5-streptavidin in solution, relative decrease of fluorescence intensity after the sample incubation, streptavidin molecular weight, and the surface area of the foil coupons as follows:

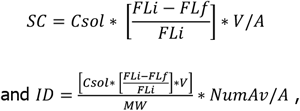

where SC is a surface concentration (ng/mm^2^), ID is an immobilization density (number/mm^2^), Csol is a concentration in solution (ng/ml), FLi if an initial fluorescence intensity, FLf is a final fluorescence intensity, V is a volume of solution (ml), MW is a molecular weight, NumAv is the Avagadro number and A is a surface area (mm^2^).

### Chandler loop assay

Stainless steel samples shaped as hollow cylinders were modified with PAB only (n=3) or with PAB, sulfo-NHS-LC-LC-biotin and avidin (n=3). The samples were inserted in appropriately sized PVC Tygon class VI tubing segments that were mounted on a Chandler loop apparatus. Each tubing segment included one control and one avidin-conjugated specimen spaced 2 cm apart. The tubing loops were filled with 10 ml of 5×10^5^/ml biotin-modified RAEC suspension in complete EGM2 medium, and the cells were re-circulated across the inserts at 25 dyn/cm^2^ and 37°C for 45 min. The inserts were then retrieved, fixed in 10% formalin overnight, stained with Hoechst 33342, cut longitudinally, flattened to expose the internal surface and examined by fluorescence microscopy. The number of fluorescent nuclei per microscopic field was counted in 3 fields per specimen.

### Direct comparison of pre-deployment and post-deployment cell seeding on stents

Stents modified with PAB, sulfo-NHS-LC-LC-biotin and avidin were assigned into two groups. Both groups of stents were then incubated with biotinylated eGFP/Luc-RAEC and expanded *ex vivo*, however the sequence of these two steps was different: cell immobilization and 12 atm balloon inflation in the pre-deployment seeding group (n=3), and 12 atm balloon inflation and cell immobilization in the post-deployment group (n=3). The external surface of stents was carefully rubbed to remove the cells attached to the external (abluminal) aspects of the struts. The stents were then extensively washed with PBS and individually placed in cell culture incubator in complete EGM2 medium for 24 hours. Luciferin (Cayman Chemical, Ann Arbor, MI) was added to the media (50 µg/ml final concentration) and the stents were examined using bioluminescence imaging (IVIS Spectrum, Waltham, MA).

### Animal experiments

All experiments involving animals were pre-approved by the Institutional Animal Care and Use Committee and were performed according to applicable federal regulations. The technique of stent deployment in the carotid arteries of Sprague-Dawley rats is described in detail in our publication[26]. Immediately after deployment of bare metal or avidin-modified stent and retrieval of the angioplasty catheter, the isolated segment of the carotid artery was flushed with PBS via the 24G cannula inserted through the arteriotomy in the external carotid artery. The cannula was then advanced to the mid-stent position and the 5×10^5^ biotin-modified or non-modified RAEC (either eGFP/Luc-expressing or non-transduced) were delivered to the stented segment and dwelled for 5 min. Cell suspension was then aspirated back in the syringe and the circulation in the common carotid artery was re-established. In the experiments assessing cell attachment to the stents, the animals (n=8) were euthanized 15 min after re-establishing carotid circulation. The stented arteries were retrieved, and formalin-fixed. The stents were then removed, cut open, flattened, stained with Hoechst 33342, and placed on microscopic slides to visualize the internal (adluminal) surface of the struts by fluorescence microscopy (Eclipse TE300, Nikon).

In the optical imaging experiments, animals (n=4) underwent bioluminescent imaging (IVIS Spectrum, Waltham, MA) after application of 100 µl of 10 µg/ml luciferin/Pluronic gel directly to the stented arterial segment[26]. After euthanasia the stented arteries were harvested and re-imaged *ex vivo*.

Finally, for the therapeutic study, rats treated with both PAB/biotin/avidin-modified stents and biotinylated RAEC (n=12), and the group treated with BMS (control; n=12) were euthanized 2 weeks after stenting. Harvested arteries were formalin-fixed, de-stented with the nitric/hydrofluoric acid formulation[27], paraffin-embedded, sectioned and stained with Verhoef/van Gieson elastic stain. Morphometric measurements were conducted on at least 5 sections for each artery and the averaged values were used for comparison between groups.

### Statistics

In all studies the chosen sample size was empirically based on prior experience with related models and experimental settings. Data are presented as mean ± standard deviation. Kolmogorov-Smirnov method was used to test for normal distribution of values within the experimental groups. Statistical analysis of differences between groups was performed using Student T-test (for 2-groups comparison) or using ANOVA (for multiple group comparison) followed by Tukey’s post-hoc test. Statistical significance was assigned with p⩽0.05.

## Results

The goal of our method for the stent surface passivation is to achieve a higher yield of EC on the adluminal stent surface after the post-deployment compared with the pre-deployment cell seeding on stents. To compare the relative number of stent-associated cells with these two seeding scenarios, biotin-modified luciferase-expressing rat aortic endothelial cells were attached to avidin-modified stents either before or after *ex vivo* expansion of the stents. Bioluminescence imaging of stents demonstrated an 11-fold higher signal emitted by the stents that were expanded after avidin immobilization but before cell attachment than the stents expanded after cell attachment (Fig. 2). The linearity between the bioluminescent signal strength and the number of luciferase-expressing cells was shown by others[28] and in our preliminary experiments.

**Fig. 2.**
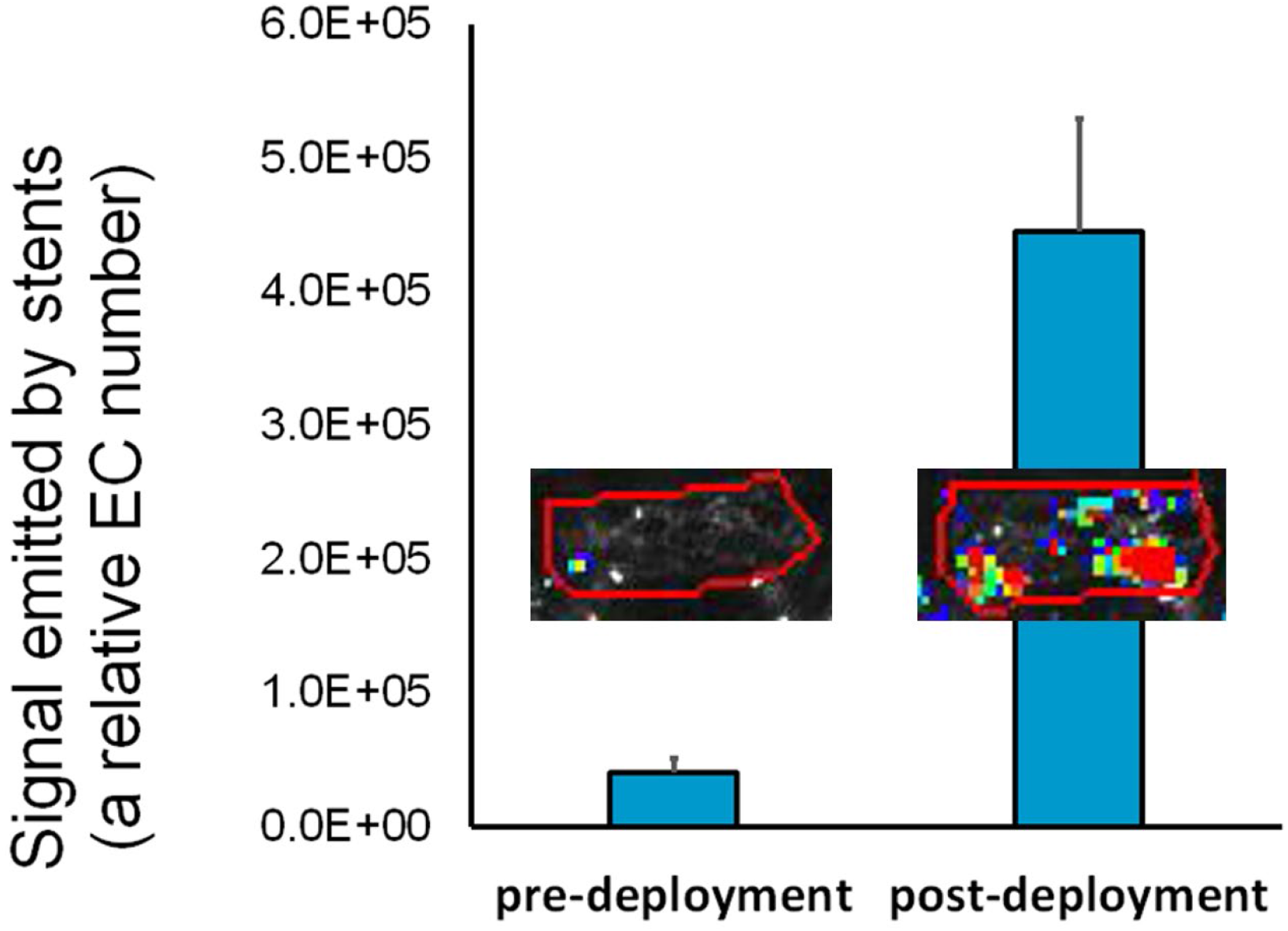
A comparison of pre-deployment and post-deployment rat endothelial cell (RAEC) seeding on stents. Luciferase-expressing biotin-modified RAEC were immobilized on avidin-modified stents either before or after *ex vivo* expansion at 12 atm. The stents were then removed from the balloons and cultured for 24 hours prior to bioluminescence imaging. The average signal strengths and the representative bioluminescence images (insets) are shown. Regions of interest are enclosed within the red line.

To assure that biotinylation of the cell surface proteins does not significantly affect cell physiology, the growth pattern of eGFP/Luc-RAEC and their expression of encoded transgenes was compared between cells treated with 4mM sulfo-NHS-LC-LC-biotin and their untreated counterparts. The cell density (Fig. 3 A-C), transgene expression (Fig. 3 A-C) and morphology (Fig. 3 B, C) did not differ significantly between the treated and non-treated groups. Biotin labeling was relatively stable despite the continuing turnover of the cell surface proteins as evidenced by staining non-modified and biotinylated RAEC with fluorescently labeled avidin 72 hours post-seeding (Fig. 3 D, E).

**Fig. 3.**
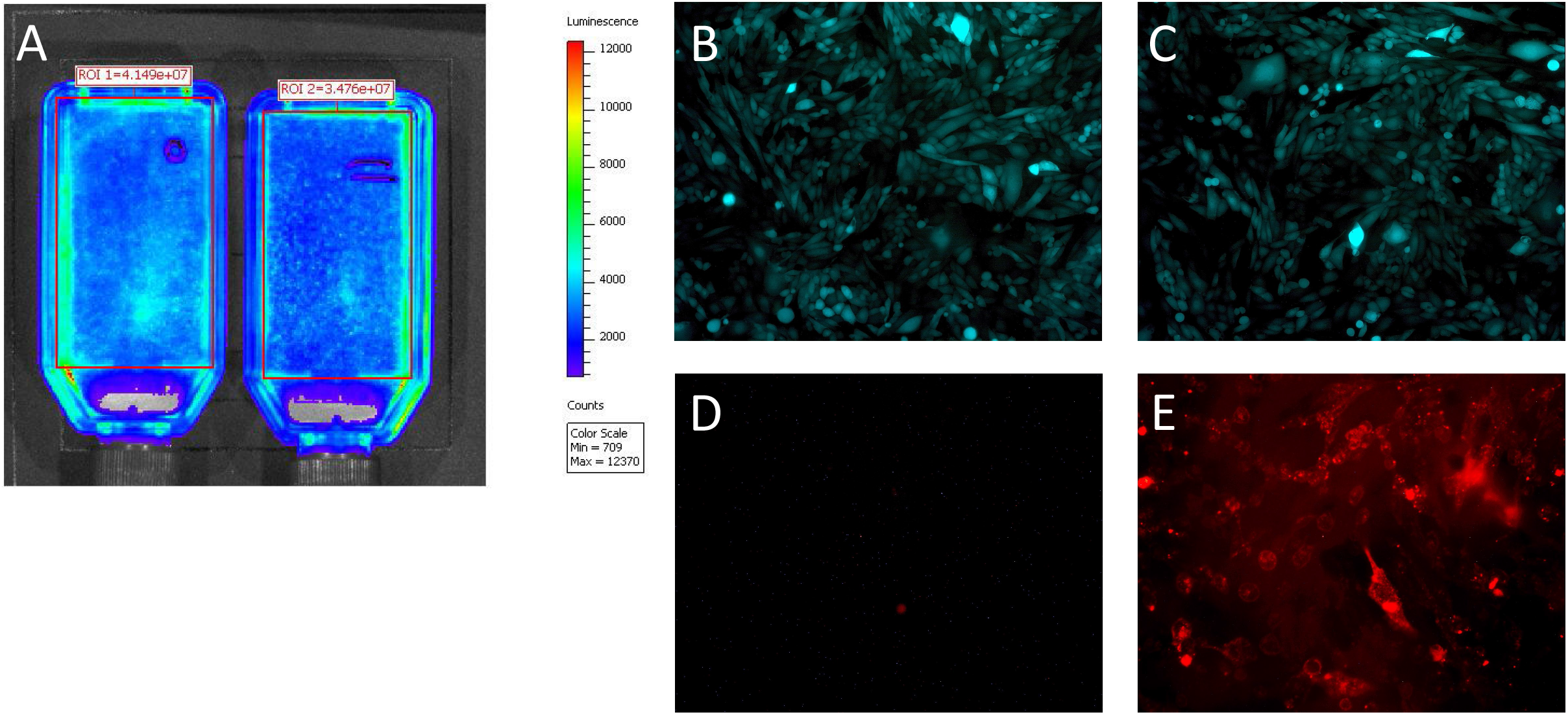
The effects of biotinylation on growth rate, morphology and transgene expression by rat endothelial cells (RAEC). RAEC expressing both luciferase and eGFP were surface-modified with sulfo-NHS-LC-LC-biotin (4mM) or were left unmodified. Equal numbers of biotinylated (A, C and E) and non-modified (A, B and D) RAEC were seeded in T-25 flasks and cultured for 72 hours. The impact of biotinylation on the RAEC growth rate and transgene expression was assessed by bioluminescence imaging (A) and fluorescence microscopy (B and C). The persistence of biotin on the surface of biotinylated cells was determined by fluorescence microscopy after treatment of cultured RAEC with Cy3.5-labeled avidin (D and E). Original magnification is 100x (B, C) and 200x (D, E).

To verify an increased avidin attachment on the biotin-primed metal surfaces and to quantify the amount of immobilized avidin, unmodified stainless steel foil samples and foil samples consecutively modified with PAB and sulfo-NHS-LC-LC-biotin were exposed to Cy3.5-labeled avidin. Depletion of fluorescent avidin in the solution with two types of metal surfaces and controls that did not include stainless steel foil specimens was assessed by fluorimetry (Fig. 4). Since the input concentration of Cy3.5-avidin and the surface area of foils were known, the surface concentration and the monolayer density of avidin immobilized via the interactions with biotin exposed on the metal surface was calculated as 4.3 ng/mm^2^, or ∼3.8×10^10^ molecules/mm^2^.

**Fig. 4.**
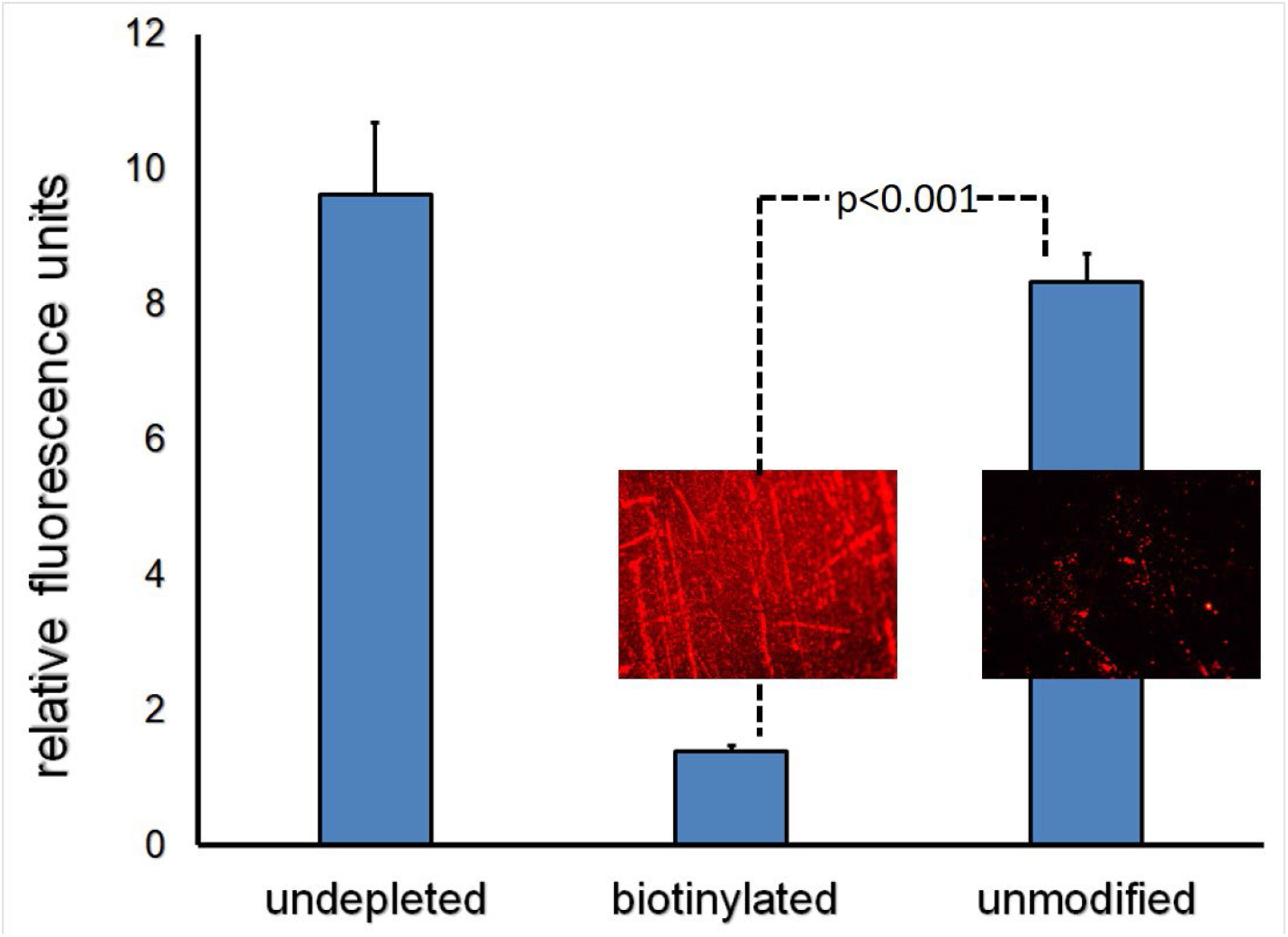
Avidin depletion assay. Unmodified stainless-steel foils (n=3) and foils consecutively modified with PAB and sulfo-NHS-LC-LC-biotin (n=3) were individually exposed to 2 ml of 1 µg/ml solution of Cy3.5-labeled avidin in 1%BSA/PBS. Control samples (n=3) did not include metal foils. Avidin attachment to the foils was then quantified by measuring Cy3.5 fluorescence of the solution depleted by the foils relative to the undepleted samples. Representative fluorescent micrograph images of unmodified and biotin-modified foils are shown as insets (200x original magnification).

Biotinylation increased cell attachment to the avidin-modified cell culture plastic more than 40-fold (Fig. 5 A, D) compared to the attachment of non-modified eGFP/Luc-RAEC (Fig. 5 C, D). This enhanced cell binding was related to the biotin/avidin interactions since biotin-modified eGFP/Luc-RAEC did not preferentially attach to the wells that was not pre-treated with avidin (Fig. 5 B, D). Importantly, biotin/avidin binding did not preclude spreading and growth of the seeded cells as evidenced by cell morphology and density after 3 days of culture (Fig. 5 E).

**Fig. 5.**
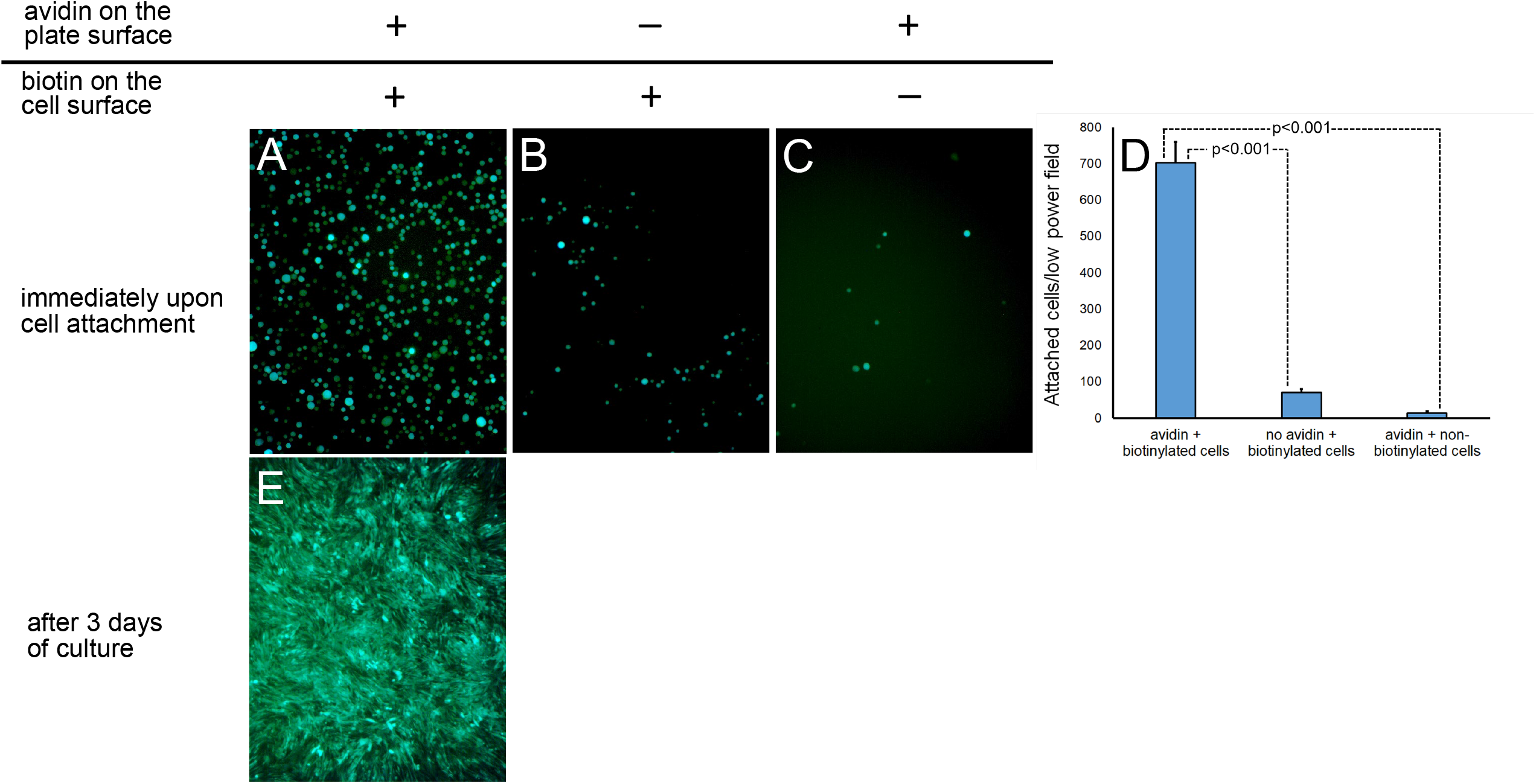
Biotinylated and control RAEC attachments to the avidin-modified and bare metal surfaces. The wells of a 96-well plate coated overnight with collagen and treated with sulfo-NHS-LC-LC-biotin were further exposed to avidin (A and C) or left untreated (B). eGFP/Luc-expressing RAEC were surface-modified with biotin (A and B) or used unmodified (C). The RAEC were added to their respective treatment wells for 15 min. The wells were imaged by fluorescence microscopy (100x). Attached GFP-positive cells were counted and the number of cells attached to the wells (n=3) under different experimental conditions was plotted (D). The cells were cultured for 72 hours and re-imaged (E).

Since the increased attachment of biotinylated cells to avidin-modified substrate under stationary conditions could not ensure robust formation of the EC monolayer under physiological flow, we exposed the avidin coated or PAB-only coated cylindrical stainless steel inserts to biotinylated RAEC recirculating through the loops of a Chandler loop apparatus and observed a 50-fold higher number of Hoechst-stained RAEC nuclei on the luminal surface of avidin-modified (Fig. 6 A, C) compared to the control inserts (Fig. 6 B, C).

**Fig. 6.**
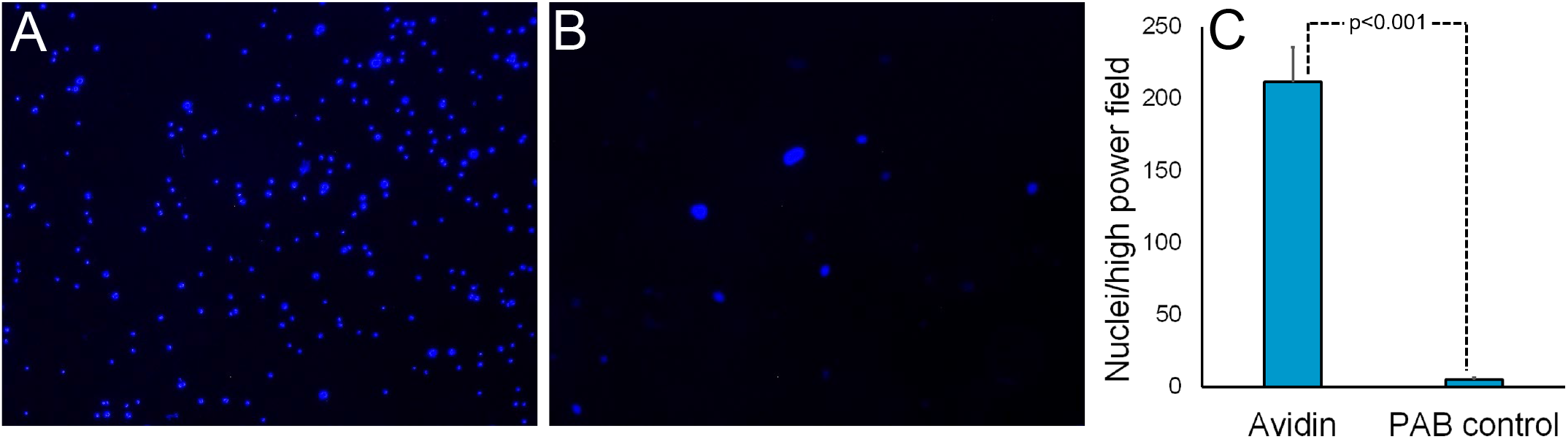
Chandler loop study. Avidin was immobilized on stainless-steel tube inserts (n=3) via PAB/sulfo-NHS-LC-LC-biotin conjugation. Control samples (n=3) were modified with PAB only. The inserts were placed into a Chandler loop apparatus filled with a suspension of biotin-modified RAEC. The cells were recirculated over the samples for 45 min (37°C, 25 dynes/cm^2^). The inserts were removed, fixed, cut to expose inside surfaces, stained with Hoechst 33342 and examined by fluorescence microscopy. Representative mages of avidin-modified sample (A) and PAB control (B) are shown along with the quantitative analysis of the images (C). Original magnification is 100x.

To validate the preferential adherence of biotinylated endothelial cells to avidin modified stents and their endurance under the flow *in vivo*, 10^5^ non-modified RAEC or 10^5^ RAEC surface-modified with biotinylation reagent, sulfo-NHS-LC-LC-biotin, were locally delivered to the isolated stented segment of rat common carotid artery immediately after stent deployment. The cells were incubated *in situ* for 5 min. The flow was then restored for 15 min, and the animals were euthanized. A fluorescence microscopy of the adluminal surface of the stents treated with non-modified (Fig. 7 A) and biotin-modified (Fig. 7 B) RAEC demonstrated significant a difference in the number of stent-associated nuclei after cell delivery and a short recirculation period (Fig. 7 C).

**Fig. 7.**
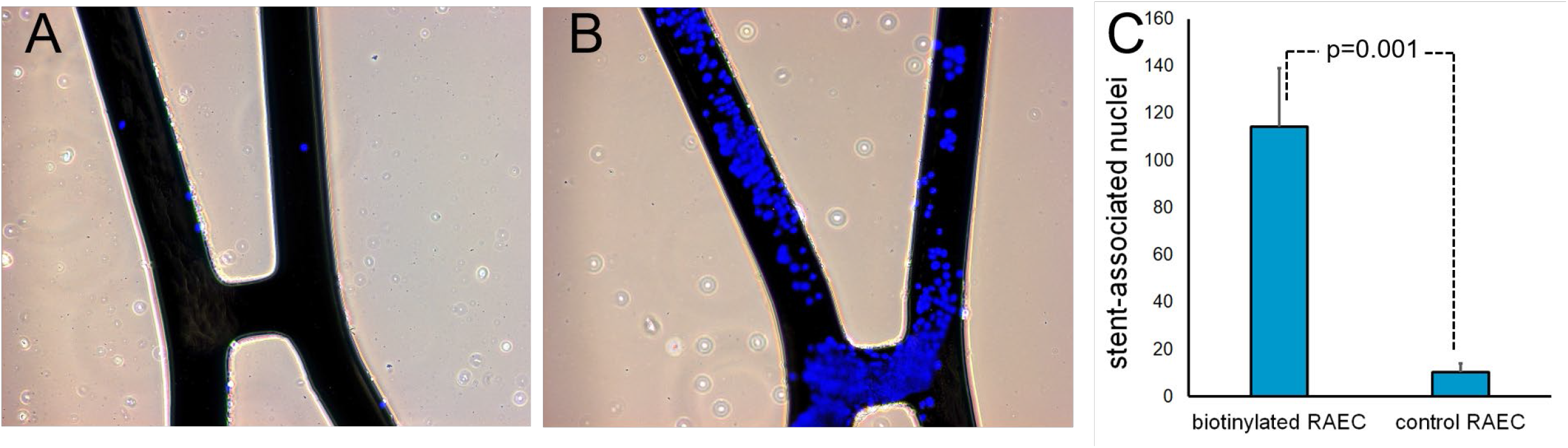
Attachment of biotinylated RAEC to avidin-modified stents in vivo. Representative images of the stents modified with avidin via PAB/biotin interactions after the common carotid artery deployment in rat model and intralumenal delivery of non-modified (A) and biotin-modified (B) RAEC. Original magnification is 100x. The number of Hoechst 33342-stained nuclei associated with the adluminal surface of struts is presented (C).

Importantly, cell immobilization via biotin/avidin tethering does not significantly affect cell physiology including the ability to express transgenes as evidenced by a 6-8 fold higher bioluminescence signal emitted from the arteries treated with avidin modified stents and delivered biotinylated eGFP/Luc-RAEC (Fig. 8 B, D) compared with the BMS-implanted arteries and similarly treated with eGFP/Luc-RAEC (Fig. 8 A, C). Finally, the post-deployment seeding of avidin-modified stents with biotinylated RAEC resulted in a 30% reduction of ISR compared to BMS-treated arteries (Fig. 8 E-G).

**Fig. 8.**
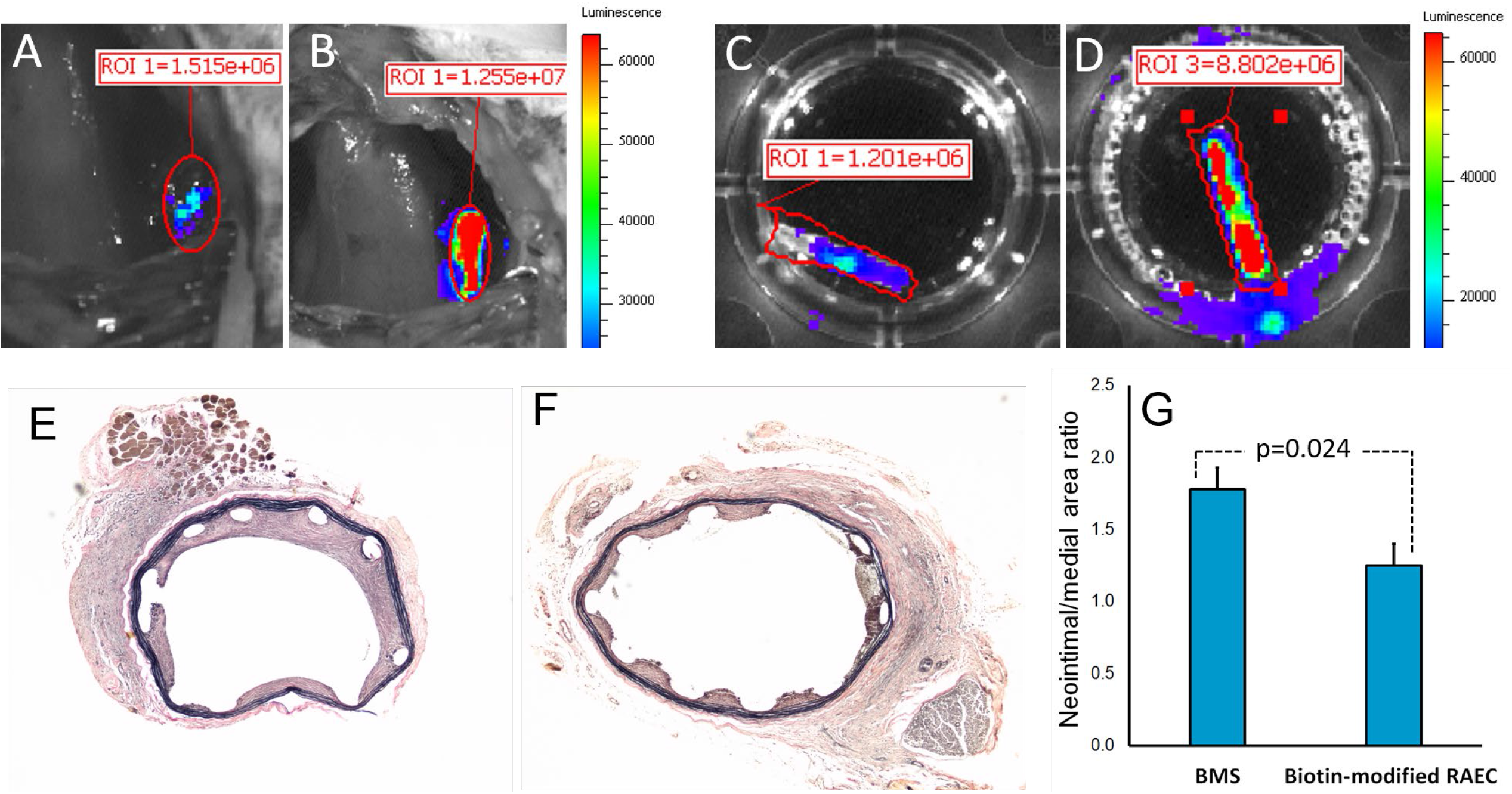
Bioluminescence imaging and therapeutic results. Bioluminescence in vivo (A, B) and ex vivo (C, D) images of stented rat carotid arteries treated with BMS (A, C) and avidin-modified stents (B, D). The stents were seeded with eGFP/Luc-RAEC immediately after deployment. Imaging was performed at 1 day post-deployment in living animals and at 2 days after deployment ex vivo. Original magnification (A-D) is 10x. Representative histological images (E, F) and morphometric analysis (G) of rat carotid arteries treated with BMS (E) and avidin-modified stents (F) and delivered biotin-modified RAEC immediately after stent deployment. E, F – original magnification is 40x.

## Discussion

Attempts to prevent ISR through accelerated re-endothelialization of stent surfaces date back to the dawn of the stent era when Dichek and Anderson[9] investigated seeding of stent struts with sheep EC, *in vitro* expansion of the stent-bound cells and their retention after stent expansion. This and subsequent [14,10,11,13,29,15] pre-deployment endothelial seeding studies fell short of clinical success mainly due to poor cell survival and retention secondary to physical stress experienced by EC during high pressure deployment. A principal novelty of our approach consists in reversing the sequence of events. Cell seeding takes place *in situ* after the stent has already been implanted. This course avoids deployment-associated cell injury and greatly increases cell retention and viability (Fig. 2). To achieve fast immobilization of EC on the stent surface we used a well-characterized affinity binding interaction between biotin and avidin. Immobilization of biotinylated EC on avidin coated surfaces was investigated in a limited manner in the context of vascular grafts and demonstrated more efficient attachment of biotin-modified cells to fibronectin and avidin coupled ePTFE grafts compared to fibronectin-only coupled grafts[30,31]. After implanting the seeded grafts in rat femoral arteries, the cell retention under flow conditions was higher in the fibronectin/avidin derivatized implants than in fibronectin-only modified counterparts[30]. However, no long-term effects of enhanced endothelialization on graft patency and neointima formation were investigated in these studies.

In parallel with development of *in vitro* stent seeding protocols, the idea of stent surface modification for promoting post-deployment attachment of blood-borne endothelial precursor cells (EPC) *in vivo* was implemented in several iterations of “pro-healing” stents [17–19,23,20,21], exemplified by the Genous™ stent that is functionalized with humanized CD34 antibody for capturing circulating EPC. While conceptually elegant, this strategy has not met with much clinical success [23], presumably due to a paucity of functional EPC in the target patient population [32]. Additionally, carbodiimide chemistry typically used to append the EPC-capturing antibodies to stent surfaces results in a random orientation of immunoglobulin molecules with a non-uniform presentation of the antigen binding epitopes[33]. In this study we investigated a new approach of achieving stent endothelialization at the time of stent implantation. To the best of our knowledge, no prior studies have investigated the concept of intra-procedural delivery of EC modified to attain rapid attachment to complementary-modified stent struts. While our studies focus on the stent coverage with cells of endothelial lineage, the overall approach is innovative in a broader sense of intra-procedural delivery and arranged immobilization of cells on the surface of implanted medical devices.

Biotinylation of RAEC did not significantly affect cell viability, proliferation and expression of reporter transgenes (Fig. 3 A-C). This finding is consistent with previously shown effects of cell surface biotinylation on the physiology of mouse mesenchymal cell lines[34]. Plasma membrane protein turnover is heterogenous and varies between 10 to 95 hours[35]. Persistence of biotin label on the plasma membrane proteins is substantiated by surface staining with fluorescent avidin 2 days after EC biotinylation and immobilization (Fig. 3 E). Prolonged presence of biotin on the cell surface may potentially be detrimental for proliferation and migration of seeded cells if the focal biotin/avidin contact points exerts sufficiently high binding force to preclude cell motility. It appears this is not the case since the attached cells can spread and grow as apparent from our data (Fig. 3 A-C and 5 E) and the literature[34,36].

Avidin depletion assay was used to determine the surface concentration and immobilization density of avidin on the biotin-modified steel surface (Fig. 4). The respective values were found to be 4.3 ng/mm^2^, or ∼3.8×10^10^ molecules/mm^2^. These amounts are consistent with previously reported surface concentration of a monolayer BSA arrangement on stainless steel[37].

Specificity of cell binding via biotin/avidin coupling in our experiments is evident from a significantly reduced number of attached RAEC when either avidin modification of the metal surface or biotin modification of cells were omitted (Fig. 5 A-D). Cell binding to substrates via integrin (Kd∼10^−5^ M)[38] is 10 orders of magnitude weaker than biotin binding to avidin (Kd 1.5×10^−15^ M). Nevertheless, EC tethering to the substrate through strong affinity forces bestowed by avidin biotin pairing does not preclude cell spreading, proliferation and migration (Fig. 5 E). Undeterred spreading and migration of endothelial cells on biotinylated substrates was previously observed[36,39,40] and explained by augmented integrin-mediated signaling[36].

Cell seeding under stationary conditions is more efficient than under flow[41]. In our Chandler loop experiments a shear stress of 25 dyn/cm^2^ allowed the attachment of biotinylated RAEC to avidin modified steel surfaces (Fig. 6). This finding indicates that biotin/avidin coupling enables stent seeding by circulating cells and expands the treatment option beyond the local delivery to regional [42] and systemic[43] administration of endothelial cells. Retention of endothelial cells immobilized on the stent struts via avidin/biotin coupling was also confirmed *in vivo* (Fig. 7).

The viability and functional competence of biotinylated RAEC attached to the stents was demonstrated by detecting bioluminescence signal emitted from the stents 1 day after stenting (Fig. 8 A, B). This signal was specifically attributed to the stented artery and not to the perivascular tissue by the *ex vivo* imaging performed 2 days after delivery (Fig. 8 C, D). The intensity of the bioluminescence signal was at least one order of magnitude higher than previously shown by us for high-gradient magnetic field mediated delivery of magnetic nanoparticle loaded EC to magnetizable stents[44,42]. Our present bioluminescence study is, however, limited by a small number of tested animals (n=2 for the group). Additionally, imaging data for delayed time points are needed to verify the survival of delivered biotinylated EC for the entire period of neointimal expansion.

The arteries of animals that received avidin-modified stents and were delivered biotinylated RAEC showed a 30% reduction of the neointima-to-media areas ratio when compared with rat carotid arteries treated with BMS (Fig. 8 E-G). This scope of restenosis inhibition is comparable with the data reported in several animal models of endothelial and EPC delivery to balloon-injured[45,46] or stented[47] arteries. The therapeutic value of this approach can be increased when the seeded cells are genetically engineered to overexpress genes with proven anti-restenotic effects, i.e. nitric oxide synthase[48,49], VEGF[16] or apoA1[50,51].

In conclusion, our proof-of-concept studies substantiate a new approach for *in-situ* stent functionalization with endothelial cells. This facile method of stent endothelialization avoids the massive cell loss on the adluminal stent surfaces with the deployment of stents seeded *in vitro* and greatly increases the efficiency of post-deployment attachment of circulating endothelial cells and EPC.

